# Neuroimaging Correlates of Altered Sense of Agency in First-Episode Schizophrenia-Spectrum Patients: A Comparative Study Across Two Sites

**DOI:** 10.1101/2025.10.10.681418

**Authors:** Karolína Volfíková, David Tomeček, Filip Španiel, Jaroslav Tintěra, Jan Rydlo, Jaroslav Hlinka

## Abstract

**Background:** Disturbances in the sense of agency are a key marker of schizophrenia and are closely linked to core symptoms such as hallucinations and delusions of influence. Neuroimaging studies have implicated cortical midline structures and the default mode network (DMN) associated with self-related processing, but replication across sites and larger samples is needed.

**Methods:** We examined neural correlates of self versus other-agency judgments in two independent cohorts of first-episode schizophrenia patients (*n* = 177) and controls (*n* = 123) recruited at separate MRI centers. During fMRI, participants performed an agency-related task. Data was analyzed using independent component analysis and statistical parametric mapping, focusing on networks associated with self-referential processing.

**Results:** Across both sites, self-agency consistently engaged the anterior and posterior cingulate cortex and the precuneus, key regions of the DMN. Schizophrenia patients exhibited significantly reduced DMN activation duri ng self-referential processing compared to controls. While site-related differences were noted in the magnitude of activation, the core pattern of DMN disruption was robust across analytic approaches and datasets.

**Conclusions:** This study replicates and extends prior findings of altered self-agency processing in schizophrenia, demonstrating consistent DMN hypo-activation in first-episode patients across two independent cohorts. These results validate the agency paradigm as a reliable probe of self-related neural mechanisms and support models of schizophrenia as a disorder of altered self-perception.

## INTRODUCTION

### Self-related processing in schizophrenia

Schizophrenia is a chronic, highly disabling, and heterogeneous mental disorder with a reported global prevalence of 0.3 to 0.6 % (1,2). It typically emerges in early adulthood and is characterized by a wide range of symptoms that affect a person’s emotions, thoughts, and behavior (3).

The heterogeneity of schizophrenia manifests in brain imaging, electrophysiology, and neurocognitive assessments (4–6). Long-term studies have shown significant variability in the course of the disorder among individuals (6), yet MR-imaging studies consistently report gray matter (GM) abnormalities in schizophrenia, particularly in the anterior cingulate cortex (ACC) and the insula (7,8). Torres et al. (9) extend these pathologies from ACC and insula to medial and inferior frontal gyri, superior temporal gyrus, medial temporal regions, and thalamus.

Several studies and meta-analyses have confirmed that the listed brain regions play a crucial role in mediating self-related processing in healthy subjects (10,11). In particular, research has demonstrated the involvement of anterior and posterior cortical midline structures (CMS), including the medial prefrontal cortex (PFC), ACC, posterior cingulate cortex (PCC), and precuneus (11). Beyond mapping of these regions, Qin and Northoff (10,12) showed that self-referential processing activates the ACC, and that this activity overlaps with resting-state activity within the default mode network (DMN). Similarly, van der Meer and colleagues (13) summarize how subdivisions of the medial PFC serve different aspects of self-reflection. Together, these findings suggest that self-related processing in healthy individuals relies on interactions within cortical midline structures, pointing to a potential neural basis for the characteristic disturbances of self-experience in schizophrenia.

### Sense of agency

The concept of the self has been explored and refined since the late 19th century. In recent neuroscience discourse, the prevailing approach divides the sense of self into two categories: the *minimal self* and the *narrative self*, as defined by Gallagher (14). The minimal self refers to an immediate, present-moment awareness of oneself as a subject of experience, unextended in time, and limited to what is accessible to immediate self-consciousness. The narrative self, also known as self-image, is, on the other hand, shaped by past experiences and future projections, constructed through the various stories we and others tell about ourselves.

Within the framework of the minimal self, Gallagher describes a phenomenon that he calls *self-agency* (14). Self-agency refers to a state in which an individual perceives themselves as the agent responsible for causing or generating one’s own actions. Gallagher explains that when a person senses that they are the source of their own actions, they are experiencing *self-agency*.

Disturbances in the sense of self are a commonly observed pathology associated with schizophrenia (15). A connection has been identified between these disruptions and the phenomenology of first-rank schizophrenia symptoms, such as auditory hallucinations, thought insertions, and delusions of influence (16,17). In many cases, pathological self-related processing precedes the onset of psychosis and is considered a core feature of schizophrenia (15,18).

### The agency experiment

Self-related experience is rarely the focus of neuroimaging studies, despite being a core marker of schizophrenia. To investigate this key symptom of the disorder, Španiel et al. designed an fMRI event-related experiment examining self-agency versus other-agency judgments (18). Using independent component analysis (ICA), Španiel et al. identified key neural networks involved in self/other-agency processing and compared activity in these regions between the two groups.

Španiel and colleagues identified three components specifically correlated with the stimulus function: parts of the default mode network (DMN) were associated with the other-agency and part of the central executive network (CEN) with the self-agency judgment. Notably, the two networks displayed antagonistic activity during the agency-related fMRI experiment.

Their results suggest that schizophrenia patients exhibit significant differences in cortical activation during the self-agency experiences in comparison to controls. Overall, patients with schizophrenia showed reduced activity in cortical midline structures.

### Replicability crisis

In recent years, concerns about the replicability of scientific research have become increasingly prominent. According to Ioannidis (19), a substantial proportion of published findings may be unreliable, primarily due to publication bias, selective reporting, and insufficient statistical power. This issue is particularly pronounced in neuroscience, where studies often suffer from small sample sizes, thereby limiting their robustness and undermining the reliability of reported results (20,21).

In this study, we aim to investigate whether the examined experimental approach is broadly effective across different sites and whether previous results can be validated and potentially expanded. We explore whether the experiment remains effective for different cohorts of participants and whether its outcomes hold across datasets collected at different sites. By addressing these questions, we aim to assess the experiment’s applicability and potential for broader use.

## METHODS AND MATERIALS

### Participants

The study sample consisted of two separate datasets, each collected at a different MRI center. The first dataset was obtained from the Institute for Clinical and Experimental Medicine in Prague (IKEM) and included data from 55 controls (CO) and 81 first-episode schizophrenia patients (SZ). The second dataset was collected at the National Institute of Mental Health in Klecany (NUDZ), comprising 68 control subjects (CO) and 96 SZ patients.

SZ patients were diagnosed according to ICD-10 criteria. The fMRI procedure was conducted during the initial stage of second-generation antipsychotic therapy. Controls were assessed using a modified version of MINI International Neuropsychiatric Interview (22) and were included in the study only if they had no lifetime history of major psychiatric disorders or a family history of psychotic disorders. Exclusion criteria for both groups included a history of seizures, significant head trauma, intellectual disability, substance dependence, or any contraindications for MRI.

Before participation, the study was explained to all participants, and written informed consent was obtained. The protocol was approved by the institutional review boards of the Prague Psychiatric Center and the National Institute of Mental Health.

In addition to sociodemographic data, clinical data were also collected, including assessments such as the PANSS scale (23) and the WHOQOL (24).

**Table 1:** Sociodemographic data, IKEM.

| | SZ ($n = 81$) | CO ($n = 55$) |
| --- | --- | --- |
| Age, years; mean (SD) | 29.4 (7.2) | 25.8 (4.3) |
| Female, no. (%) | 40 (49.4) | 31 (56.4) |
| Male, no. (%) | 41 (50.6) | 24 (43.6) |

**Table 2:** Sociodemographic data, NUDZ.

| | SZ ($n = 96$) | CO ($n = 68$) |
| --- | --- | --- |
| Age, years; mean (SD) | 28.8 (7.3) | 31.9 (8.2) |
| Female, no. (%) | 38 (39.6) | 41 (60.3) |
| Male, no. (%) | 58 (60.4) | 27 (39.7) |

### fMRI Acquisition

#### IKEM

The acquisition of MRI data collected at IKEM was performed on a 3 Tesla Siemens Trio scanner equipped with a standard 12-channel head coil. For anatomical reference, structural 3-dimensional (3D) images were obtained using the T1-weighted (T1w) magnetization-prepared rapid gradient echo (MPRAGE) sequence with the following parameters: repetition time (TR) of 2300 ms, echo time (TE) 4.6 ms, flip angle 10°, voxel size of 1 × 1 × 1 *mm*^3^, field of view (FOV) 256 *mm* × 256 *mm*, matrix size 256 × 256, 224 sagittal slices. Functional images were obtained using the T2*-weighted (T2*w) gradient echo-planar imaging (GR-EPI) sequence sensitive to the blood oxygenation level-dependent (BOLD) signal with following parameters: repetition time (TR) of 2000 ms, echo time (TE) 30 ms, flip angle 90°, voxel size of 3 × 3 × 3 *mm*^3^, field of view (FOV) 192 *mm* × 192 *mm*, matrix size 64 × 64, each volume with 30 axial slices (slice order: sequential decreasing), 240 volumes in total.

#### NUDZ

The acquisition of MRI data collected at NUDZ was obtained using a 3 Tesla Siemens Prisma scanner equipped with a standard 64-channel / 20-channel head coil. For anatomical reference, structural 3D images were obtained using the T1w magnetization-prepared rapid gradient echo (MPRAGE) sequence with the following parameters: TR of 2400 ms, TE 2.3 ms, flip angle 8°, voxel size of 0.7 × 0.7 × 0.7 *mm*^3^, FOV 224 *mm* × 224 *mm*, matrix size 320 × 320, 240 sagittal slices. Functional images were obtained using the T2*w gradient echo-planar imaging (GR-EPI) sequence sensitive to the BOLD signal with following parameters: TR of 2000 ms, TE 30 ms, flip angle 90°, voxel size of 3 × 3 × 3 *mm*^3^, FOV 192 *mm* × 192 *mm*, matrix size 64 × 64, each volume with 30 axial slices (slice order: alternating increasing), 240 volumes in total.

### Task and Design

The self-agency judgment in subjects was assessed using an experimental approach first introduced by Španiel et al. (18). During the fMRI session, participants were presented with stimuli in the form of a scene, as shown in Figure 1. The stimuli were projected onto a mirror attached to the head coil via a screen positioned at the head end of the scanner bore. Participants interacted with the stimuli using an MRI-compatible joystick, which moved the cursor on the screen.

**Figure 1.**
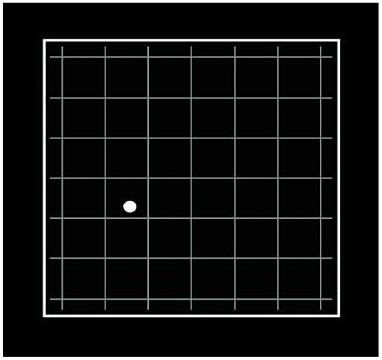
Representative screenshot of the stimuli projected during the agency experiment in the functional magnetic resonance imaging (fMRI) (18).

Participants were instructed to move the cursor based on whether they perceived its movement as fully corresponding to their joystick inputs or not. They were informed beforehand that, during the experiment, they would occasionally observe cursor movements that appeared to be controlled by the experimenter from outside the scanner. In such cases, if they subjectively interpreted the movement as influenced by the experimenter, participants were instructed to keep moving the cursor inside the central square. Conversely, if they perceived the cursor’s movement as fully aligned with their joystick inputs, they were to guide the cursor to the outer corridor of the screen. Participants were not allowed any cessation of the movement.

The experiment was structured as alternating blocks of other-agency (OA) and self-agency (SA). During OA blocks, software-based random angular distortions were introduced to the cursor’s movement, independent of the participant’s joystick input. While the speed of the cursor movement remained proportional to the velocity of joystick movements, angular distortions in the polar coordinate system were applied to the direction of the movement. During SA blocks, the cursor movement corresponded precisely to the joystick inputs made by the participants.

The task consisted of 12 OA blocks and 12 SA blocks, each lasting 20 seconds. There were no visual-feedback distortions between the blocks. Participants were blinded to the sequentiality and length of the experimental blocks. Debriefing revealed no evidence for recognition of either regularity or artificiality of this approach. All participants also underwent a 3-minute training period to ensure they fully understood the task.

### Preprocessing

The preprocessing steps were similar across both datasets (IKEM, NUDZ), when applied separately. Initially, the fMRI images were converted from DICOM to NIFTI format using the dcm2niix tool (25). The preprocessing pipeline then followed standard procedures including bias field correction, realignment and unwarping, slice-timing correction, direct segmentation, spatial normalization into standard sterotactic space (EPI template; Montreal Neurologic Institute, MNI-152), and smoothing with a Gaussian kernel (8 × 8 × 8 *mm*^3^ full width at half maximum). This pipeline is referred to as the default preprocessing pipeline for volume-based analyses (direct normalization to MNIspace) in the CONN toolbox (26,27).

Additionally, we performed screening of the fMRI data to ensure adequate field-of-view (FOV) coverage and to identify potential motion artifacts. In cases of insufficient FOV coverage or excessive image distortion caused by motion artifacts, the data of this subject were excluded from the study.

### Task-related ICA

The analysis of task-related brain activity was conducted using group spatial Independent Component Analysis (ICA). The analysis was performed on each dataset separately. Initially, two standard Principal Components Analysis (PCA) reduction steps were performed on the fMRI data. The ICA was then computed using the GIFT toolbox (28,29). The number of independent components (ICs) required was pre-defined as 35, based on the previous study (18).

We based the selection of relevant independent components on several steps. First, ICs needed to demonstrate stability as assessed by the ICASSO algorithm implemented in the GIFT toolbox. ICASSO performs multiple ICA iterations with varying initial conditions to evaluate the robustness of the estimated ICs. For our analysis, we configured the ICASSO to run 20 ICA iterations. ICs with a stability index below the commonly used threshold of 0.9 were excluded.

To identify task-related ICs, we calculated the correlation coefficient between each component’s mean time series and the stimulus function (*z >* ±3.18). The stimulus function was generated as alternating blocks of zeros (for OA) and ones (for SA) convolved with the hemodynamic response function. ICs that did not exhibit a significant correlation with the task were excluded.

The final criterion for IC selection involved a between-group comparison of the components’ time series. We conducted a two-sample t-test on the beta weights (regression coefficients) derived from regression analyses on the components (*p* < 0.05, FWE corrected). ICs that did not show significant differences between the groups were removed.

The resulting ICs were depicted as spatial maps thresholded at *z >* 3. The identification of artifacts was subject to visual assessment based on the article by Kiviniemi et al. (30). Anatomical labels were resolved using the Talairach Daemon Atlas (31,32).

### Voxel-wise statistical analysis

The next approach to analyzing fMRI data from the agency experiment involved voxel-wise statistical analysis using SPM12 (33,34). This analysis was conducted separately for each dataset. The stimulus function was generated in a manner similar to the approach described in the previous section.

First, we generated individual first-level contrast images (p < 0.05, FWE corrected) for each condition (OA/SA). Subsequently, a one-sample t-test was performed to produce within-group activation maps for both the CO and SZ groups. Next, we conducted a two-sample t-test at the whole brain level (p < 0.05, FWE corrected, minimal cluster size > 20 voxels) to compare the activation patterns between the CO and SZ groups. To exclude the influence of age, we considered it a confounding factor, regressing it out during the two-sample t-tests.

## RESULTS

### Task-related ICA

#### IKEM

Independent Component Analysis (ICA) was applied separately to each dataset, estimating 35 components for each. In the IKEM dataset, six components were labeled unreliable by the ICASSO algorithm and excluded. Additional selection steps, based on the correlation with the stimulus function and the two-sample t-test, led to the exclusion of several other components. To summarize, ICA identified six ICs (C6, C13, C14, C18, C19, C26) significantly related to the task and showed significant differences between the two groups (CO, SZ), as shown in Figure 2.

**Figure 2.**
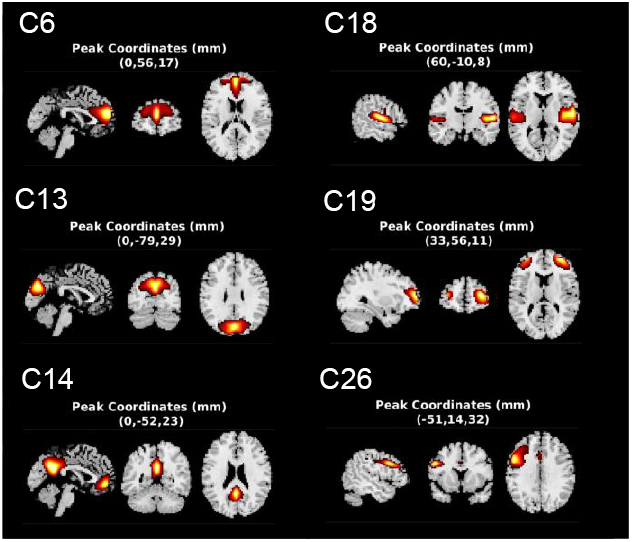
Task-related independent components (ICs) of the IKEM dataset, identified using the GIFT toolbox (26,27), are shown as spatial maps. ICs were found based on regression analysis of the mean IC time series against the stimulus function. Connectivity z-scores are represented on a red-to-yellow scale.

C6 primarily mapped to the anterior cingulate cortex (BA 24, 32) and the anterior prefrontal cortex (PFC) (BA 10), both corresponding to the anterior segment of the default mode network (aDMN). C13 corresponded to the visual cortex (BA 17, 18, 19), while C14 was predominantly localized in the posterior cingulate cortex and precuneus (BA 23, 31), both playing a pivotal role in the DMN (pDMN), while appearing also in the anterior cingulate (BA 24, 32). C18 corresponded to the auditory cortex (BA 41). C19 corresponded to the superior, middle, and inferior frontal gyri bilaterally (BA 10, 46) and was identified as part of the central-executive network (CEN). Finally, C26 was identified as part of the fronto-parietal network (BA 9, 44).

#### NUDZ

For the NUDZ dataset, ICA found four task-related components (C12, C17, C18, C21), as shown in Figure 3.

**Figure 3.**
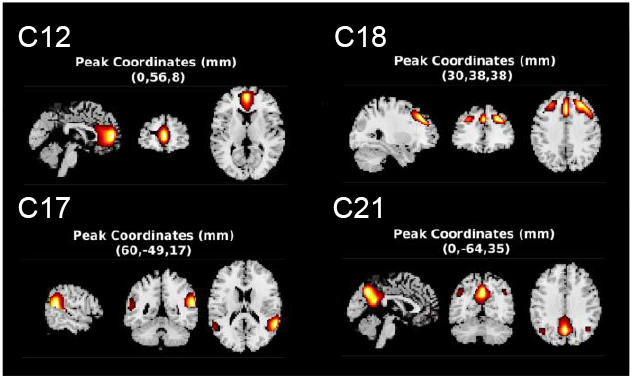
Task-related independent components (ICs) of the NUDZ dataset, identified using the GIFT toolbox (26,27), are shown as spatial maps. ICs were found based on regression analysis of the mean IC time series against the stimulus function. Connectivity z-scores are represented on a red-to-yellow scale.

Component C12 corresponded to the anterior cingulate cortex and anterior PFC (BA 10, 24, 32). C17 was also identified as a part of the DMN due to its correspondence with the temporo-parietal junction (BA 32). C18 corresponded to the dorsomedial PFC (BA 8, 9) and C21 to the posterior segment of the DMN, including the posterior cingulate cortex and precuneus (BA 23, 31).

### Voxel-wise analysis

Using a one-sample t-test, we examined whether each group exhibited significant brain activity during each task condition separately. The results are depicted as spatial maps in Figure 4 for the IKEM dataset and Figure 5 for the NUDZ dataset, respectively.

**Figure 4.**
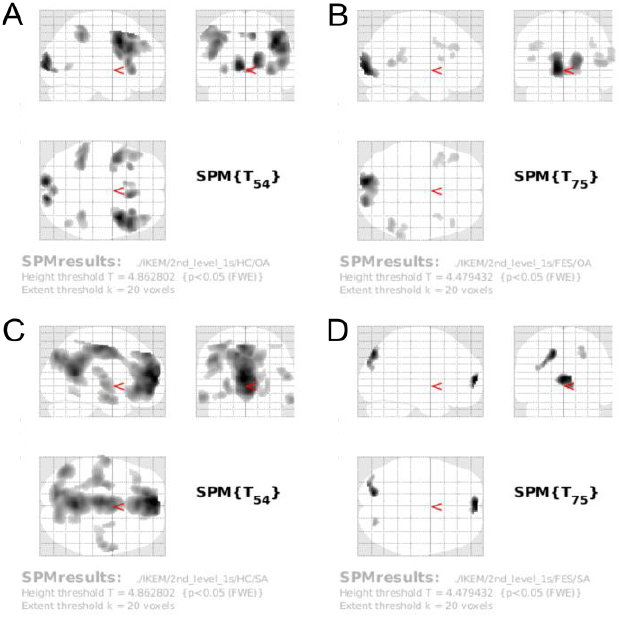
Spatial t-maps showing the results of a whole-brain within-group t-test conducted on the IKEM dataset. p < 0.05, FWE corrected, min. cluster size > 20. Maps show clusters in which there were significant activations within the group (controls, CO/schizophrenia patients, SZ) during a given condition - other-agency (OA) / self-agency (SA): A) OA, CO; B) OA, SZ; C) SA, CO; D) SA, SZ.

**Figure 5.**
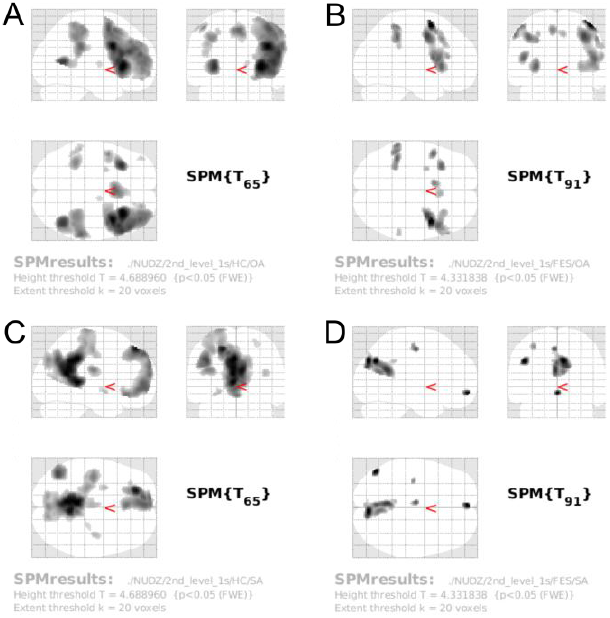
Spatial t-maps showing the results of a whole-brain within group t-test conducted on the NUDZ dataset. p < 0.05, FWE corrected, min. cluster size > 20. Maps show clusters in which there were significant activations within the group (controls, CO / first-episode-schizophrenia patients, SZ) during a given condition-other-agency (OA) / self-agency (SA): A) OA, CO; B) OA, SZ; C) SA, CO; D) SA, SZ

In both datasets, during the self-agency condition, significant activation was observed in the ACC, PCC, precuneus, and both the anterior and dorsolateral PFC in the control group. In SZ patients, we found significant activity only in the anterior PFC and visual cortex in the case of the IKEM dataset. In the NUDZ dataset, however, activation was more prominent in the visuo-motor cortex rather than the visual cortex.

During the other-agency condition, both datasets showed prominent activation in the premotor cortex in the case of controls. Among SZ patients, activation was observed in the visual cortex in the IKEM dataset and in the premotor cortex in the NUDZ dataset.

#### Group comparison in IKEM

Between-group differences were tested during the SA condition. Since the results for the OA condition are equivalent to the SA condition, only with switched test setting (*CO > SZ* for OA condition is equivalent to *SZ > CO* for SA condition), in Figure 6, we show the results only for SA. The age of the participants was considered a confounding factor and regressed out.

**Figure 6.**
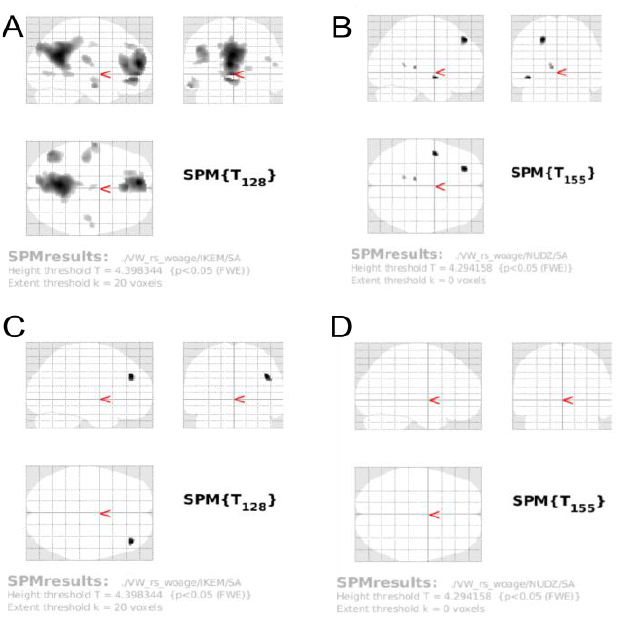
Spatial t-maps showing the results of a whole-brain between-group (controls CO, first-episode schizophrenia patients SZ) analysis during the self-agency (SA) condition. p < 0.05, FWE corrected, min. cluster size > 20. Maps show clusters in which: A) CO>SZ in the IKEM dataset; B) CO>SZ in the NUDZ dataset; C) SZ > CO in the IKEM dataset; D) SZ > CO in the NUDZ dataset; where “>” symbolizes stronger activation resp. deactivation.

In the IKEM dataset (A, C in Figure 6), there were two major areas found in which the CO group showed significantly greater activations in comparison to the SZ group (*p <* 0.05, FWE corrected, minimal cluster size *>* 20 voxels). The areas comprise the anterior and posterior cingulate cortex, the anterior PFC, and the precuneus.

One cluster was found in which the SZ group exhibited significantly greater activation than the CO group (*p <* 0.05, FWE corrected, minimal cluster size *>* 20 voxels). The cluster was located in the left dorsolateral PFC (BA 9).

#### Group comparison in NUDZ

In the case of the NUDZ dataset (B, D in Figure 6), there were no clusters found in which the SZ group would exhibit significantly greater activations than the CO group (*p <* 0.05, FWE corrected, minimal cluster size *>* 20 voxels), during the SA condition. For the opposite test setting, *CO > SZ*, there were a few clusters in which the condition was met (*p <* 0.05, FWE corrected, minimal cluster size *>* 20 voxels). The clusters were localized in the right dorso-lateral PFC (BA 9) and the right supramarginal gyrus (BA 40).

## DISCUSSION

The results of this study support the findings of Španiel et al. (18), while extending them through the use of a larger sample and data collected across two independent sites. One dataset was obtained from the Institute for Clinical and Experimental Medicine (IKEM), representing an expanded version of the sample used in the Španiel et al. study. The second dataset was independently acquired from the National Institute of Mental Health (NUDZ) - and did not include any participants from the previous research. Across both datasets, we identified several brain regions that were consistently activated during self-and other-agency judgment tasks. These findings are consistent with prior literature (10,11), which associates self-related processing with activation in cortical midline structures.

Using the independent component analysis (ICA), we identified brain regions associated with the experimental task. Notably, there was an overlap between the significant ICA components from the previous study and those found in both tested datasets. These regions included the anterior and posterior cingulate cortex, as well as the precuneus. Consistent with the earlier study, we classified these regions as part of the default mode network (DMN) and observed similar activity patterns. The BOLD signal in these areas was positively correlated with the self-agency condition and negatively with the other-agency condition.

Our analysis also discovered other brain regions associated with the experimental task but not reported in the study of Španiel and colleagues. Namely, in the case of the dataset from IKEM, we found the visual and auditory cortex to be also significantly engaged during the task, both positively correlated with the self-agency condition and negatively with the other-agency condition. In the case of the dataset from NUDZ, there was a significant activation in the dorsomedial prefrontal cortex (PFC), which was positively correlated with the other-agency condition and negatively with the self-agency condition.

The results suggest that the agency experiment arouses activity in brain regions associated with self-oriented perception. Additionally, ICA identified regions that were significant only within a particular dataset.

Similar conclusions emerged from the voxel-wise analysis. The results show that in controls, ACC, PCC, precuneus, and parts of the PFC were significantly activated during the self-agency condition. During the other-agency condition, controls primarily exhibited activity in the premotor cortex.

In contrast, SZ patients did not show significant activations in the PCC or precuneus during self-agency. Additionally, activation in the region of the premotor cortex, during other-agency, was notably weaker in SZ patients.

The two-sample t-test revealed that the differences between the two groups are mainly due to the weaker activation of ACC and PCC in the group of SZ patients during the self-agency condition.

The group of patients exhibited decreased DMN resp. CEN activation resp. deactivation during the task, compared to controls. This finding was supported by two different analytical approaches – independent component analysis and voxel-wise analysis – and was also consistent across both study sites.

However, our analyses also revealed notable differences between the two examined datasets. These differences primarily stemmed from variations in the degree of activation and deactivation in specific brain regions within each group, rather than from the involvement of different brain regions. Overall, the control and SZ groups appeared more similar in terms of task-related BOLD signal in the NUDZ dataset than in the IKEM dataset. A post-hoc analysis indicated that the SZ groups did not differ significantly between the two datasets. Instead, the primary difference was observed in the control groups, with the IKEM controls exhibiting stronger activations in certain clusters compared to the NUDZ controls (see Figure 8 in the Supplement).

We had already considered age as a confounding factor, but the differences between the samples remained even after regressing it out. To determine the cause of the different effects, we looked at other potential confounding factors that could contribute if they were not equally distributed across groups. We found a significant difference in the distribution of the PANSS scale (*U* = 5537.5, *p <*.001), where the mean score for all patients in the IKEM dataset was significantly higher than in the NUDZ group, though the primary difference stemmed from the control group. This aligns with the results of the WHOQOL scale comparison, when we examined the parameter indicating the overall health (G1.2) as perceived by the participant - we found a significant difference between the SZ groups (*U* = 4340.5, *p <*.01). The SZ group from the IKEM dataset had overall higher scores on the G1.2 parameter in the WHOQOL scale compared to the SZ group from NUDZ. The difference was notable between the groups of controls, too (*U* = 1328, *p <*.01). Additionally, we examined whether task compliance indices or other demographic variables could provide further insights, but no significant differences were found between the datasets. For further details, see Table 3 in the Supplement.

Considering all the results and additional exploratory analyses, we conclude that the observed differences are likely due to other site-related effects. Between-site variability appeared mainly in effect magnitude rather than spatial pattern, likely reflecting differences in acquisition parameters and sample demographics. Nevertheless, we confirmed that the self-agency experiment activates regions of the anterior and posterior cingulate cortex and the precuneus, which are associated with self-related processing. Importantly, in connection with this result, we found that schizophrenia patients exhibit significantly reduced activation of the default mode network during self-referential processing, supporting a model of altered self-perception in schizophrenia.

In summary, first-episode schizophrenia is associated with reduced DMN engagement during self-agency judgments across two independent sites. These results highlight disrupted self-related processing as an early neural marker of schizophrenia and support further validation of the agency task as a translational probe.

## ACKNOWLEDMENTS

This study was co-funded by the European Union (BRADY, No. CZ.02.01.01/00/22_008/0004643; BRAINSCAPE, No. CZ.02.01.01/00/23_020/0008560) and Ministry of Health of the Czech Republic (Grant nr. NU21-08-00432).

Preliminary results were presented at the OHBM 2025 conference in Brisbane as a poster number 1061. A previous version of this article was published as a preprint:

Volfíková K, Tomeček D, Španiel F, Tintěra J, Rydlo J, Hlinka J (2025): Altered Sense of Agency in First-Episode Schizophrenia Patients: A Comparative Study Across Two Sites. *bioRxiv* 2025.10.10.681418

I would like to thank my colleagues Ing. Katarzyna Kazimierczak, Ph.D. for her assistance with the identification of brain regions and Ing. Anna Pidnebesna, Ph.D. for valuable insights during the development of this work.

## REFERENCES

1. Charlson FJ, Ferrari AJ, Santomauro DF, Diminic S, Stockings E, Scott JG, et al. (2018): Global Epidemiology and Burden of Schizophrenia: Findings From the Global Burden of Disease Study 2016. Schizophr Bull 44: 1195–1203.

2. Saha S, Chant D, Welham J, McGrath J (2005): A Systematic Review of the Prevalence of Schizophrenia ((S. E. Hyman, editor)). PLoS Med 2: e141.

3. American Psychiatric Association (Ed.) (1998): Diagnostic and Statistical Manual of Mental Disorders: DSM-IV; Includes ICD-9-CM Codes Effective 1. Oct. 96, 4. ed., 7. print. Washington, DC.

4. Liu Z, Palaniyappan L, Wu X, Zhang K, Du J, Zhao Q, et al. (2021): Resolving heterogeneity in schizophrenia through a novel systems approach to brain structure: individualized structural covariance network analysis. Mol Psychiatry 26: 7719– 7731.

5. Luo C, Pi X, Hu N, Wang X, Xiao Y, Li S, et al. (2021): Subtypes of schizophrenia identified by multi-omic measures associated with dysregulated immune function. Mol Psychiatry 26: 6926–6936.

6. Jiang Y, Luo C, Wang J, Palaniyappan L, Chang X, Xiang S, et al. (2024): Neurostructural subgroup in 4291 individuals with schizophrenia identified using the subtype and stage inference algorithm. Nat Commun 15: 5996.

7. Glahn DC, Laird AR, Ellison-Wright I, Thelen SM, Robinson JL, Lancaster JL, et al. (2008): Meta-Analysis of Gray Matter Anomalies in Schizophrenia: Application of Anatomic Likelihood Estimation and Network Analysis. Biol Psychiatry 64: 774–781.

8. Bora E, Fornito A, Radua J, Walterfang M, Seal M, Wood SJ, et al. (2011): Neuroanatomical abnormalities in schizophrenia: A multimodal voxelwise meta-analysis and meta-regression analysis. Schizophr Res 127: 46–57.

9. Torres US, Duran FLS, Schaufelberger MS, Crippa JAS, Louzã MR, Sallet PC, et al. (2016): Patterns of regional gray matter loss at different stages of schizophrenia: A multisite, cross-sectional VBM study in first-episode and chronic illness. NeuroImage Clin 12: 1–15.

10. Northoff G, Heinzel A, De Greck M, Bermpohl F, Dobrowolny H, Panksepp J (2006): Self-referential processing in our brain—A meta-analysis of imaging studies on the self. NeuroImage 31: 440–457.

11. Murray RJ, Schaer M, Debbané M (2012): Degrees of separation: A quantitative neuroimaging meta-analysis investigating self-specificity and shared neural activation between self-and other-reflection. Neurosci Biobehav Rev 36: 1043–1059.

12. Qin P, Northoff G (2011): How is our self related to midline regions and the default-mode network? NeuroImage 57: 1221–1233.

13. Van Der Meer L, Costafreda S, Aleman A, David AS (2010): Self-reflection and the brain: A theoretical review and meta-analysis of neuroimaging studies with implications for schizophrenia. Neurosci Biobehav Rev 34: 935–946.

14. Gallagher S (2000): Philosophical conceptions of the self: implications for cognitive science. Trends Cogn Sci 4: 14– 21.

15. Jeannerod M (2009): The sense of agency and its disturbances in schizophrenia: a reappraisal. Exp Brain Res 192: 527–532.

16. Lindner A, Thier P, Kircher TTJ, Haarmeier T, Leube DT (2005): Disorders of Agency in Schizophrenia Correlate with an Inability to Compensate for the Sensory Consequences of Actions. Curr Biol 15: 1119–1124.

17. Waters F, Woodward T, Allen P, Aleman A, Sommer I (2012): Self-recognition Deficits in Schizophrenia Patients With Auditory Hallucinations: A Meta-analysis of the Literature. Schizophr Bull 38: 741–750.

18. Spaniel F, Tintera J, Rydlo J, Ibrahim I, Kasparek T, Horacek J, et al. (2016): Altered Neural Correlate of the Self-Agency Experience in First-Episode Schizophrenia-Spectrum Patients: An fMRI Study. Schizophr Bull 42: 916–925.

19. Ioannidis JPA (2005): Why Most Published Research Findings Are False. PLoS Med 2: e124.

20. Button KS, Ioannidis JPA, Mokrysz C, Nosek BA, Flint J, Robinson ESJ, Munafò MR (2013): Power failure: why small sample size undermines the reliability of neuroscience. Nat Rev Neurosci 14: 365–376.

21. Szucs D, Ioannidis JPA (2017): Empirical assessment of published effect sizes and power in the recent cognitive neuroscience and psychology literature ((E.-J. Wagenmakers, editor)). PLOS Biol 15: e2000797.

22. Sheehan DV (n.d.): The Mini-International Neuropsychiatric Interview (M.I.N.I.): The Development and Validation of a Structured Diagnostic Psychiatric Interview for DSM-IV and ICD-10.

23. Kay SR, Fiszbein A, Opler LA (1987): The Positive and Negative Syndom Scale (PANSS) for Schizophrenia. Schizophr Bull 13: 261–276.

24. The Whoqol Group (1998): Development of the World Health Organization WHOQOL-BREF Quality of Life Assessment. Psychol Med 28: 551–558.

25. Li X, Morgan PS, Ashburner J, Smith J, Rorden C (2016): The first step for neuroimaging data analysis: DICOM to NIfTI conversion. J Neurosci Methods 264: 47–56.

26. Nieto-Castanon A (2020): Handbook of Functional Connectivity Magnetic Resonance Imaging Methods in CONN. Hilbert Press.

27. Whitfield-Gabrieli S, Nieto-Castanon A (2012): Conn: A Functional Connectivity Toolbox for Correlated and Anticorrelated Brain Networks. Brain Connect 2: 125–141.

28. Calhoun VD, Adali T, Pearlson GD, Pekar JJ (2001): A method for making group inferences from functional MRI data using independent component analysis. 10.1002/hbm.1048

29. Du Y, Fu Z, Sui J, Gao S, Xing Y, Lin D, et al. (2020): NeuroMark: An automated and adaptive ICA based pipeline to identify reproducible fMRI markers of brain disorders. NeuroImage Clin 28: 102375.

30. Kiviniemi V, Starck T, Remes J, Long X, Nikkinen J, Haapea M, et al. (2009): Functional segmentation of the brain cortex using high model order group PICA. Hum Brain Mapp 30: 3865–3886.

31. Lancaster JL, Woldorff MG, Parsons LM, Liotti M, Freitas CS, Rainey L, et al. (2000): Automated Talairach Atlas labels for functional brain mapping. Hum Brain Mapp 10: 120–131.

32. Lancaster J l., Rainey L h., Summerlin J l., Freitas C s., Fox P t., Evans A c., et al. (1997): Automated labeling of the human brain: A preliminary report on the development and evaluation of a forward-transform method. Hum Brain Mapp 5: 238–242.

33. Neuroimaging B members & collaborations of the WCFH (n.d.): SPM, Statistical Parametric Mapping, version 12. Retrieved from https://www.fil.ion.ucl.ac.uk/spm/

34. Ashburner J, Barnes G (2014): SPM12 manual. Wellcome Trust Centre for Neuroimaging, London, UK.

