## Supplementary material for "Neuroimaging Correlates of Altered Sense of Agency in First-Episode Schizophrenia-Spectrum Patients: A Comparative Study Across Two Sites"

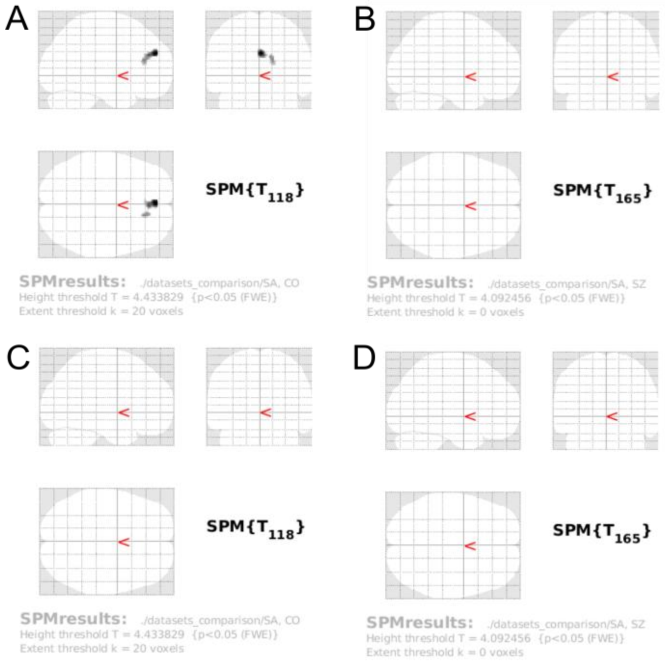


Figure 7: Spatial t-maps showing the results of a whole-brain between-sites (IKEM/NUDZ) analysis during the self-agency (SA) condition. $p < 0.05$, FWE corrected, min. cluster size > 20. Maps show clusters in which: A) IKEM > NUDZ in the C group; B) IKEM > NUDZ in the SZ group; C) NUDZ > IKEM in the C group; D) NUDZ > IKEM in the SZ group; where ">" symbolizes stronger activation resp. deactivation.

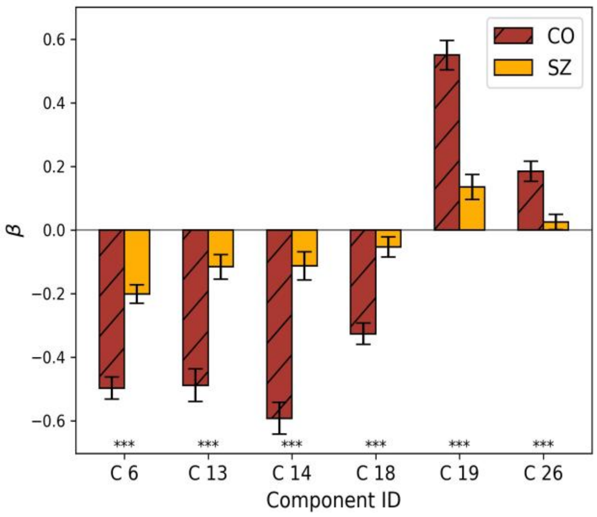


Figure 9: Results of statistical comparison of regression coefficients obtained from ICA analysis between the group of controls and the group of SZ patients in the IKEM dataset. The comparison was done using two-sample t-test conducted on the mean regression coefficient values for each component separately. The significance of the result is indicated by the sign $*$, where $***$ correspond to $p < .001$.

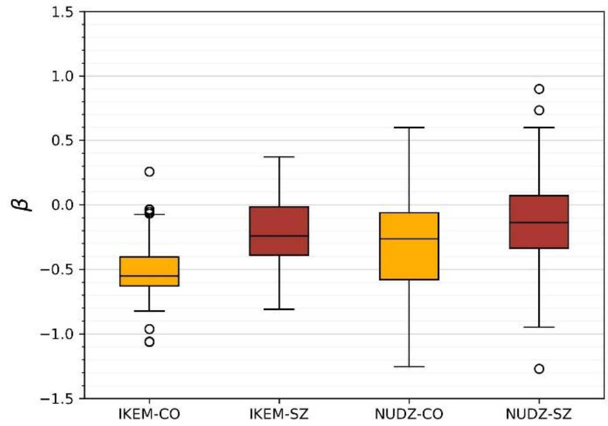


Figure 8: Distribution of regression (β) coefficients for the region of anterior DMN identified by ICA across four groups: control group from IKEM (IKEM-CO), patients group from IKEM (IKEM-SZ), control group from NUDZ (NUDZ-CO), and patients group from NUDZ (NUDZ-SZ). Each box-plot represents the distribution of β coefficients within each group, illustrating between-group differences in the brain region's activation

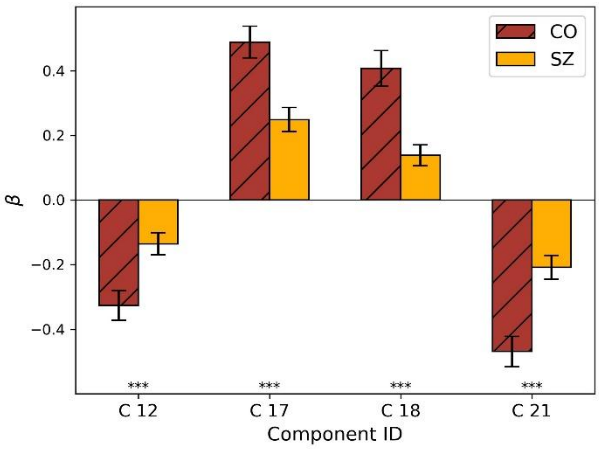


Figure 10: Results of statistical comparison of regression coefficients obtained from ICA analysis between the group of controls and the group of SZ patients in the NUDZ dataset. The comparison was done using two-sample t-test conducted on the mean regression coefficient values for each component separately. The significance of the result is indicated by the sign $*$, where $***$ correspond to $p < .001$.

| Parameter | tested groups | U | p | sign. |
|---|---|---|---|---|
| Age | $CO_I$ - $CO_N$ | 1068 | $1.00E^{-4}$ | *** |
| | $SZ_I$ - $SZ_N$ | 3583 | 0.784 | |
| | $CO_I$ - $SZ_I$ | 1457 | 0.003 | ** |
| | $CO_N$ - $SZ_N$ | 3590 | 0.051 | |
| PANSS | $SZ_I$ - $SZ_N$ | 5538 | $7.72E^{-11}$ | *** |
| QOL-H | $CO_I$ - $CO_N$ | 1328 | 0.004 | ** |
| | $SZ_I$ - $SZ_N$ | 4341 | 0.004 | ** |
| | $CO_I$ - $SZ_I$ | 640 | $7.68E^{-13}$ | *** |
| | $CO_N$ - $SZ_N$ | 2264 | 0.003 | ** |
| CI | $CO_I$ - $CO_N$ | 1872 | 0.158 | |
| | $SZ_I$ - $SZ_N$ | 3016 | 0.743 | |
| | $CO_I$ - $SZ_I$ | 3075 | $1.87E^{-6}$ | *** |
| | $CO_N$ - $SZ_N$ | 2982 | 0.003 | ** |

Table 3: Mann-Whitney U test results for examined demographical and clinical parameters: age, PANSS scale (global score), the health parameter of WHO-QOL, and compliance index (CI). Indexes I and N indicate the dataset to which the group belongs (IKEM, resp. NUDZ).
